# Single-Molecule FRET-Tracking of InlB-Activated MET Receptors in Living Cells

**DOI:** 10.1101/2025.05.06.652421

**Authors:** Yunqing Li, Marina S. Dietz, Hans-Dieter Barth, Hartmut H. Niemann, Mike Heilemann

## Abstract

The activation of transmembrane receptors through the binding of external ligands initiates information transfer across the cell membrane. Understanding these processes requires observations in living cells. Given the heterogeneity and lack of synchronization of such events, single-molecule experiments are required to resolve distinct sub-populations. Here, single-molecule FRET microscopy and single-particle tracking are combined to track the ligand-induced dimerization and activation of the MET receptor tyrosine kinase in the plasma membrane of living cells. First, using fluorophore-labeled variants of the MET ligand internalin B (InlB), the lifetime of a ligand-activated dimeric (MET:InlB)_2_ receptor complex is determined to be around 1 second. Next, diffusion coefficients of monomeric and dimeric MET:InlB complexes are extracted from single-molecule FRET trajectories, revealing an approximately 1.6-fold slower diffusion of the dimeric receptor compared to the monomeric receptor, accompanied by spatially restricted motion. The combination of single-molecule FRET and single-particle tracking provides essential biophysical parameters of membrane receptor activation in living cells.

## Introduction

The transmembrane receptor MET, also known as hepatocyte growth factor receptor (HGFR), belongs to the class of receptor tyrosine kinases (RTKs) and plays a central role in regulating key cellular functions, including proliferation, migration, morphogenesis, and tissue regeneration.^[1–3]^ Dysregulation of MET is associated with various diseases, including cancer.^[4–6]^ MET is activated upon binding of its physiological ligand, hepatocyte growth factor/scatter factor (HGF/SF),^[7,8]^ its natural isoform NK1,^[9,10]^ and the bacterial ligand InlB.^[11,12]^ Ligand binding promotes the formation of a 2:2 complex, consisting of two MET receptors and two ligands, which promotes cross-phosphorylation of MET within the complex and the subsequent initiation of downstream signaling cascades.^[13]^ MET signaling is regulated by receptor internalization and degradation.^[14–16]^

InlB plays a critical role in bacterial pathogenesis by exploiting host cell signaling machinery to facilitate bacterial entry.^[12,17,18]^ *Listeria monocytogenes*, a Gram-positive pathogen responsible for listeriosis, expresses InlB on its surface to mediate invasion into nonphagocytic cells.^[19]^ Through high-affinity interaction with MET, InlB hijacks host signaling pathways that ultimately support bacterial internalization and systemic infection.^[12,20–23]^ Therefore, a complete understanding of the dynamics and mechanisms of MET activation via InlB is important. The activity and structure of the MET:InlB dimer were studied in detail with x-ray crystallography, biochemical tools, single-molecule microscopy, and molecular dynamics simulation.^[24–29]^ However, until now, the dynamics of the MET:InlB dimer were only studied indirectly using single-particle tracking,^[25,30]^ whereas information on the stability, i.e., lifetime, of the (MET:InlB)_2_ dimer was so far inaccessible.

The lifetime of ligand-induced RTK dimers is a key determinant of signaling duration before receptor internalization or dissociation. This lifetime is regulated by interactions with ligands, co-receptors, other cellular structures (lipids, actin cytoskeleton), and downstream signaling proteins.^[31–36]^ The stability of the dimer influences the recruitment of adaptor proteins to the phosphorylated intracellular tyrosine residues generated upon dimerization, thereby modulating downstream signaling pathways.^[16,37]^ For example, prolonged dimerization of EGFR has been shown to enhance proliferative signaling.^[38,39]^ Consequently, measuring both the lifetime and diffusion properties of RTK dimers in living cells is key information to describe receptor activation kinetics and spatial dynamics within the native membrane environment. It also serves as a reference measure for the development of drugs that aim to destabilize receptor dimers or weaken receptor activation in a disease context. Because of the technical complexity of accessing this information in living cells, comprehensive studies of these parameters simultaneously remain exceedingly rare.

In this study, we determine the diffusion dynamics of InlB-bound MET monomers and dimers in living cells by combining single-molecule Förster resonance energy transfer (smFRET) and single-particle tracking (SPT). In addition, we measure the dimer lifetime and lateral mobility of MET receptors in the plasma membrane. Our results provide quantitative information on the stability and dynamics of MET:InlB dimers, complementing the biophysical model of receptor activation.

## Material and methods

### Passivation of coverslips with PLL-PEG-RGD

Glass surfaces were passivated with RGD-grafted poly-L-lysine-graft-(polyethylene glycol) (PLL-PEG-RGD) to simultaneously minimize nonspecific binding and promote cell attachment. For live-cell measurements, coverslips were sonicated in isopropanol for 20 min at 35 °C using an ultrasonic bath (S30H Elmasonic, Elma electronic GmbH, Germany). They were then rinsed three times with deionized water (ddH_2_O) and dried with nitrogen gas. Plasma cleaning was performed for 10 min using nitrogen at 80% power and approximately 0.3 mbar pressure. A 10 µL aliquot of 0.8 mg/mL PLL-PEG-RGD solution was applied between two coverslips and incubated for 1.5 h at room temperature in a 10 cm culture dish. After incubation, coverslips were gently separated, rinsed with ddH_2_O, dried under nitrogen, and transferred to 6-well plates. Coverslips were stored under argon, sealed with Parafilm, and kept at −20 °C for up to two weeks before use.^[30]^ For fixed cell measurements, 8-well chambers (SARSTEDT AG & Co. KG, Nümbrecht, Germany) were passivated with PLL-PEG-RGD according to Li et al.^[29]^

### Cell culture

The human osteosarcoma cell line U-2 OS (CLS Cell Lines Service GmbH, Eppelheim, Germany) was cultivated in high glucose Dulbecco’s modified Eagle medium/nutrient mixture F-12 (DMEM/F12, Gibco, Life Technologies, Waltham, MA, USA), supplemented with 1% GlutaMAX (Gibco, Life Technologies) and 10% fetal bovine serum (FBS) (Gibco, Life Technologies) at 37 °C with 5% CO_2_ in an automatic CO_2_ incubator (Model C 150, Binder GmbH, Tuttlingen, Germany) and passaged every 3 to 4 days.

For live cell imaging experiments, 2 mL of a suspension containing 3 ×10^4^ U-2 OS cells in complete growth medium supplemented with 1 unit/mL penicillin and 1 µg/mL streptomycin (Gibco, Life Technologies) was seeded per well in a 6-well plate (Greiner, Bio-One International GmbH, Kremsmünster, Österreich), which has been supplemented with PLL-PEG-RGD-coated glass coverslips with a diameter of 25 mm (VWR International GmbH). Cells were allowed to adhere and grow for three days prior to measurement.

For smFRET in fixed cells, U-2 OS cells were seeded in 8-well chambers (SARSTEDT AG & Co. KG, Nümbrecht, Germany) passivated with PLL-PEG-RGD. 300 µL cell suspension per well with 1.5 × 10^4^ cells supplemented with 1 unit/mL penicillin and 1 µg/mL streptomycin (Gibco, Life Technologies) was seeded for 2 to 3 days to adhere and grow before measurement.

### Sample preparation

For ligand stimulation, H and T cysteine variants (K64C, K280C) of InlB were site-specifically labeled with Cy3B and ATTO 647N using maleimide chemistry for thiol-specific labeling, as previously described^[29]^.

Prior to live-cell imaging, coverslips with U-2 OS cells were mounted into custom-built holders, and 500 µL of pre-warmed live-cell imaging solution (Gibco) was added. Cells were incubated at room temperature for 10 min to reduce temperature-induced variability. Afterward, an oxygen scavenging buffer was added, consisting of 0.009 U/µL glucose oxidase from *Aspergillus niger* (Type VII), 594 U/mL catalase from bovine liver, 0.083 M glucose, and 1 mM Trolox (all from Sigma-Aldrich, St. Louis, MO, USA). 30 nM of each Cy3B- and ATTO 647N-labeled InlB_321_ variants (T-T or H-T) were added to ensure stoichiometric labeling of MET by InlB. The InlB-containing buffer was incubated with the cells for 5 min before the measurement.

Fixed-cell smFRET experiments were performed as previously described.^[29]^ Briefly, cells were incubated with 5 nM of each InlB_321_ variant for 15 min at 37 °C and subsequently fixed for 15 min with 4% formaldehyde (Thermo Scientific) and 0.01% glutaraldehyde (Sigma-Aldrich) in 0.4 M sucrose and 1× PBS. For measurements, a freshly prepared oxygen-scavenging buffer (see above) was added.

### smFRET acquisition

Single-molecule imaging was performed using a custom-built TIRF microscope. Cells were measured at room temperature for 30 min. Two lasers, 637 nm (140 mW OBIS) and 561 nm (200 mW Sapphire) from Coherent Inc. (Santa Clara, CA, USA), were used for excitation. The laser beams were collinearly aligned via a dichroic mirror (H 568 LPXR superflat, AHF Analysentechnik AG, Tübingen, Germany) and modulated through an acousto-optical tunable filter (AOTF; AOTFnC-400.650-TN, AA Opto-Electronic, Orsay, France), which allowed for rapid alternating excitation. The switching frequency of the laser excitation was regulated by two digital counters and analog output boards (NI PCI-6602 and NI PCI-6713, National Instruments, Austin, TX, USA). After passing through the AOTF, the laser beams were coupled into a single-mode optical fiber (P5-460AR-2, Thorlabs) via a fiber collimator (PAF-X-7-A, Thorlabs, Dachau, Germany) and re-collimated to a 2 mm beam diameter upon exiting the fiber using a collimation unit (60FC-0-RGBV11-47, Schäfter & Kirchhoff, Hamburg, Germany). The collimated beams were directed onto a dual-axis galvo mirror system (GVS012/M, Thorlabs), which enabled precise control of the illumination mode between wide-field, stationary TIRF, circular TIRF, or highly inclined laminated optical sheet (HILO) illumination via a custom, Python-based control script. The excitation light was delivered into an inverted microscope (IX-71, Olympus Deutschland GmbH, Hamburg, Germany) and focused onto the back focal plane of the objective (UPlanXApo, 100x, NA 1.45, Olympus Deutschland GmbH) using a pair of telescope lenses (AC255-050-A-ML and AC508-100-A-ML, Thorlabs). Excitation and emission paths were spectrally filtered by a set of optical filters and dichroic mirrors (Dual Line Clean-up ZET561/640x, Dual Line rejection band ZET 561/640, Dual Line beam splitter zt561/640rpc, AHF Analysentechnik AG). Z-axis stability during acquisition was maintained using a nosepiece stage (IX2-NPS, Olympus Deutschland GmbH), minimizing focus drift over time. Fluorescence emission was collected by the same objective and directed to the detection pathway via the dichroic mirror. For dual-color imaging, an image splitter (Optosplit II, Cairn Research Ltd, UK) was employed, equipped with a beam splitter and two bandpass filters (H643 LPXR, 590/20 BrightLine HC, 679/41 BrightLine HC, AHF Analysentechnik AG). Both emission channels were recorded simultaneously on an EMCCD camera (iXon Ultra X-10971, Andor Technology Ltd, Belfast, UK), yielding a final optical magnification of 100× and an effective pixel size of 159 nm.

The live-cell measurements followed the smFRET recovery after photobleaching (smFRET-RAP) method ^[40]^. Specifically, the overlapping signals from the basal membrane of the cells were first photobleached 5 to 10 min using 561 nm (67.7 W/cm^2^) and 637 nm (552.1 W/cm^2^) laser illumination in TIRF mode. After allowing signal recovery from the apical and lateral membranes for 1 min without illumination, 4000 frames were acquired using 561 nm excitation (19.8 W/cm^2^) for FRET observation. Control measurements consisting of 500 frames were acquired before and after the photobleaching process to assess photobleaching and signal recovery, using 561 nm (19.8 W/cm^2^) and 637 nm (76.5 W/cm^2^) laser excitation. A transmitted light image was recorded after each measurement to confirm cell integrity. The acquisition was controlled using the µManager software^[41]^ using 40 ms integration time, 200 EM gain, 3 times preamp gain, 17 MHz readout rate, and activated frame transfer for a region of interest (ROI) of 512 ×256 px. For channel alignment, 100-frame image sequences of 100 nm TetraSpeck™ microspheres were acquired using 561 nm and 637 nm laser excitation at the start of each measurement day.

For smFRET measurements in fixed cells with alternating laser excitation, 1000 frames were acquired with an exposure time of 100 ms, activated frame transfer, an EM gain of 150, a preamp gain of 3x, a readout rate of 17 MHz, and an image size of 512 × 256 pixels. A transmitted light image was taken per measurement. The excitation lasers (561 nm: 47 W/cm^2^, 637 nm: 172 W/cm^2^) were alternated with a frequency of 10 Hz.

### Data analysis

The recorded field of view was separated into donor and acceptor channels using ROIs identified from 100 nm TetraSpeck™ calibration measurements in iSMS,^[42]^ applying the autoalign ROIs tool with default settings. The donor and acceptor channels were then tracked separately using u-track^[43]^ with the following parameters: localization using Gaussian Mixture-model fitting with data-driven PSF radius derived in 10 iterations, 2D tracking with 3 frames gap closure time, and 20 frames as the minimum length of a trajectory. The trajectories detected by u-track were further processed using the smCellFRET software^[40]^ to identify those smFRET trajectories that showed either an increase in donor intensity upon disappearance of the acceptor or a decrease in donor intensity upon an increase in acceptor intensity.

The donor trajectory is concatenated with the nearest matching donor trajectory within a 250 nm search radius at the position where the acceptor signal disappeared. After selecting the smFRET trajectories, the FRET intensity traces were summarized and exported using SPARTAN.^[44]^ The FRET efficiency was calculated based on the equations introduced by Li et al.^[29]^

Besides the intensities of donor and acceptor, the coordinates of smFRET trajectories were extracted using custom Matlab and Python scripts. Each trajectory was segmented into two parts: the fraction that is FRET-active and the donor-only fraction following the disappearance of the acceptor signal. The assignment process consisted of 3 steps: (1) Repeated localizations at the same position of donor and acceptor channels were merged to one localization; (2) Donor coordinates that spatially overlapped with the coordinates of the acceptor were identified as part of the FRET trajectory; (3) Donor coordinates with no matching acceptor signal were assigned to the donor-only trajectory segment. Trajectory characteristics, including diffusion coefficients determined by a global mean squared displacement (MSD) fit for the first four displacement points, dynamic localization precision, and type of motion, were extracted from the MSD plot according to the description in Harwardt et al.^[30]^ The lifetimes of the FRET trajectories were collected by analyzing the length of the FRET traces. The jump angles were calculated as the direction difference between individual steps within each segment using a custom Python script.^[45]^ Visualization of the results was done using OriginPro (version 2023, OriginLab Corporation, Northampton, MA, USA).

smFRET measurements in fixed cells were analyzed with iSMS. smFRET data were deposited in the EMBL BioImaging Archive (https://www.ebi.ac.uk/biostudies/bioimages/studies/S-BIAD1347).

## Results

### smFRET of (MET:InlB_321_)_2_ in live cells using smFRET-RAP

We determined the dynamics of (MET:InlB_321_)_2_ dimers from an available single-molecule FRET data set recorded in living cells that was previously used to determine absolute FRET efficiencies for the dimeric complex.^[29]^ Conceptually, two InlB_321_ variants, each with a unique accessible cysteine at either position 64 (termed H for head) or position 280 (termed T for tail) (**Figure 1A**), were fluorescently labeled with either Cy3B or ATTO 647N (**Table S1**).^[29]^ Accessible volume simulation of the crystal structure of the (MET:InlB_321_)_2_ dimer^[24]^ yielded expected FRET efficiencies between donor and acceptor of 0.620 (T-T),0.567 (H-T/T-H), and 0.018 (H-H).^[29]^ Single-molecule FRET data in living cells were recorded using fluorescence bleaching and recovery (smFRET-RAP)^[40]^ (**Figure 1B**). Using high-intensity excitation of both donor and acceptor fluorophores and TIRF illumination, fluorophore-labeled InlB_321_ bound to MET at the basal membrane or adsorbed to the glass surface was photobleached. Following this, the cells were maintained in the dark to allow MET-bound InlB_321_ to diffuse from the lateral and apical membranes back to the basal membrane, restoring a low-density signal that is suitable for single-molecule analysis. Next, smFRET video files were acquired using donor excitation and simultaneous detection of donor and acceptor fluorescence (see Methods). Only smFRET intensity traces showing anti-correlated intensity of donor and acceptor emission and one-step photobleaching were accepted for FRET efficiency calculations (**Figure S1**). Distinct populations with FRET efficiencies of 0.90 ± 0.05 for the T-T pair and 0.55 ± 0.09 for the H-T pair were found (**Figure 1C**), in good agreement with data obtained from fixed cells,^[29]^ yet broader FRET efficiency distributions were observed (**Figure S2A**).

**Figure 1:**
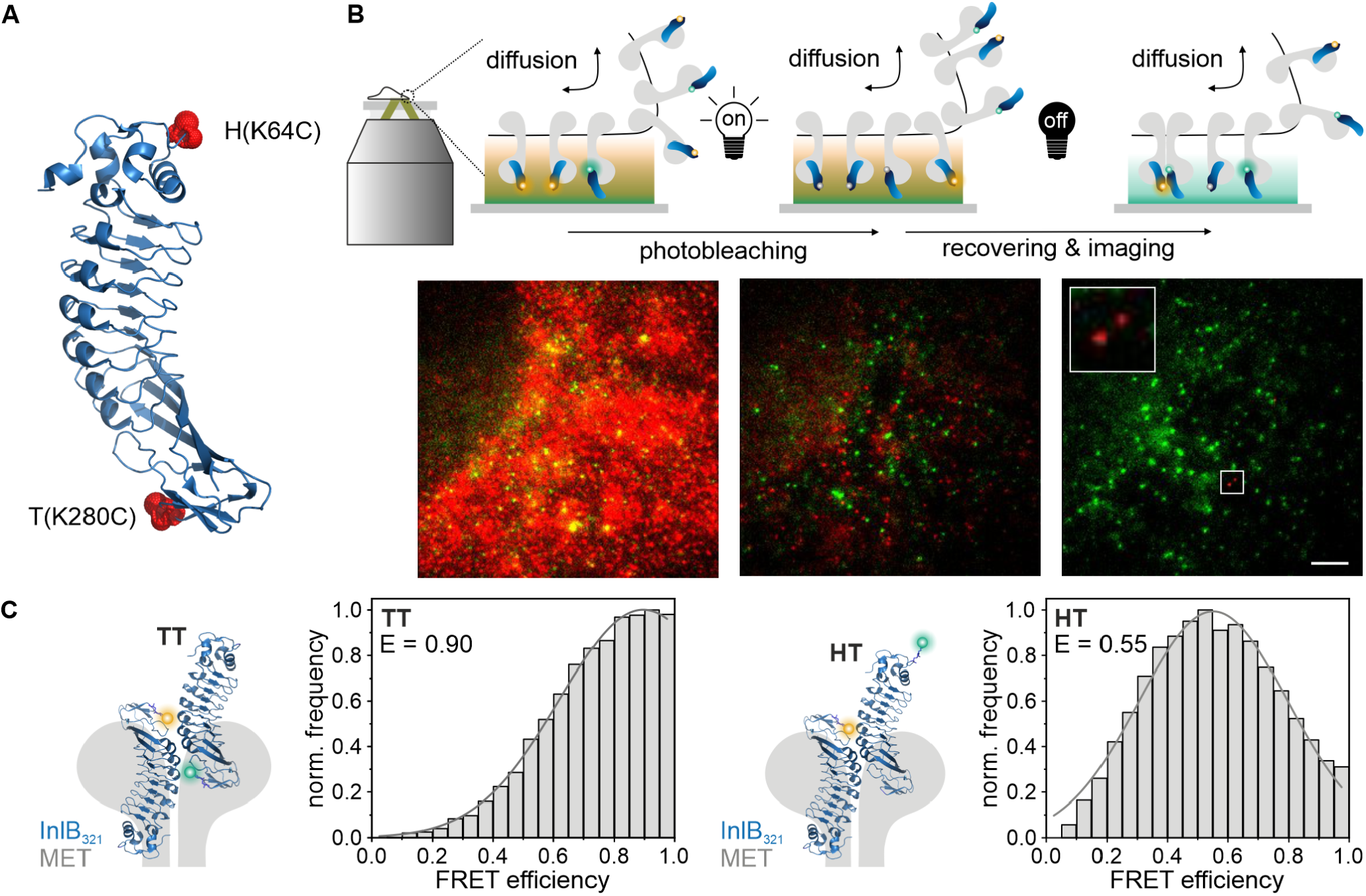
smFRET-RAP experiments of (MET:InlB)2 dimers in living cells. **A**) crystal structure of InlB321 (PDB 1H6T). The H (K64C) and T (K280C) mutations are highlighted in red. The two variants were labeled with either Cy3B or ATTO 647N, respectively. **B**) Schematic of smFRET-RAP (top) with example image before and after photobleaching, and during the single-molecule FRET measurement (bottom). Scale bar 5 μm. **C**) Live-cell smFRET efficiencies of (MET:InlB321)2 dimers labeled with Cy3B-T-InlB321 and ATTO 647N-T-InlB321 (left) or with Cy3B-H-InlB321 and ATTO 647N-T-InlB321 (right). The crystal structure of InlB is adapted from PDB: 2UZY.

### smFRET reports the lifetime of the dimeric (MET:InlB)_2_ complex in living cells

To determine the lifetime (association time) of dimeric (MET:InlB)_2_ complexes in living cells, we reconstructed single-molecule trajectories and measured the duration of the FRET signal in these trajectories. First, a single-molecule FRET event is identified at the moment a signal in the acceptor emission channel is observed upon donor excitation (**Figure 2A**). Next, the single-molecule FRET signal is followed over time and a trajectory is reconstructed, including donor and acceptor fluorescence signals until both signals disappear (**Figure 2B; SI Movies 1-4**). The temporal length of the FRET segment of the trajectory was used to calculate the apparent lifetime of the dimeric (MET:InlB)_2_ complex and was found to be 1.13 ± 0.06 s (T-T) and 0.80 ± 0.03 s (H-T) for the two FRET pairs, respectively (**Figure 2C**). In order to assess whether photobleaching is the major determinant of the trajectory length, we analyzed single-molecule FRET data of (MET:InlB)_2_ complexes in fixed cells. The total lifetime of single-molecule FRET was found to be about 15-fold longer than in living cells (**Figure S2B**), indicating that the FRET lifetime in living cells is mainly dominated by the lifetime of the (MET:InlB)_2_ complex. Furthermore, we observed that in about 30% of all FRET trajectories, the donor fluorescence emission persists after the acceptor signal disappears (**Figure S3A, E**).

**Figure 2:**
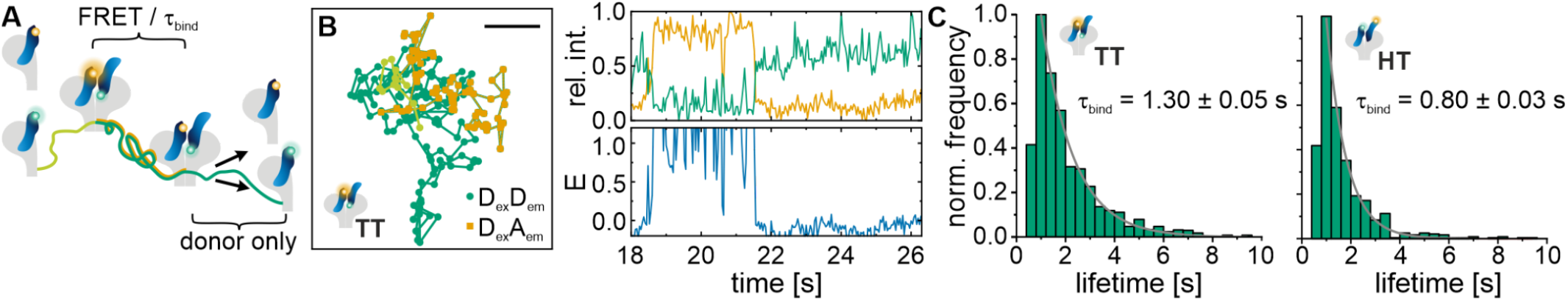
Lifetime of the (MET:InlB)2 dimer. **A**) Scheme of a live-cell smFRET trajectory. Single-donor fluorescence that turns into a FRET signal is the starting point for a single-molecule trajectory, which is then recorded until both the donor and the acceptor disappear. **B**) Exemplary smFRET trajectory. The left panel shows the positions of the acceptor and donor, respectively, before (light green) and during FRET (orange), and after the acceptor signal disappeared (green). In the right panels, the corresponding intensities (donor in green, acceptor in orange) and FRET efficiencies over time are shown. Scale bar 500 nm. **C**) Histograms of the duration of the FRET segments in single-molecule trajectory deliver (MET:InlB)2 dimer lifetimes for InlB T-T and H-T variants. The histograms were fitted with a single exponential decay. N = 27 cells (T-T) and 24 cells (H-T), 3 independent experiments.

### Diffusion properties of the dimeric (MET:InlB)_2_ and monomeric (MET:InlB) are distinct

Next, the diffusion coefficients and types were extracted from single-particle trajectories exhibiting FRET. Both segments of the trajectories were analyzed separately, the first one where FRET was detected, and the second one that followed the loss of acceptor signal and showed donor fluorescence only (**Figure 3**). For MET labeled with InlB T–T variants, the mean diffusion coefficient of FRET segments was 0.066 ± 0.044 µm^2^/s, while donor-only segments exhibited a higher mean diffusion coefficient of 0.109 ± 0.068 µm^2^/s. In cells where the MET receptor was labelled with H–T variants, the mean diffusion coefficients were 0.056 ± 0.038 µm^2^/s for FRET trajectories and 0.093 ± 0.053 µm^2^/s for donor-only trajectories. The distribution of diffusion coefficients showed a clear separation between FRET and donor-only segments, with the latter consistently displaying higher mobility (**Figure 3A**). Analysis of the types of motion revealed that 44% (T-T) and 50% (H-T) of FRET trajectories were classified as restricted diffusion, while the remaining 56% (T-T) and 50% (H-T) showed free diffusion. In contrast, in the donor-only segments, a smaller proportion (39% (T–T) and 33% (H–T)) showed restricted diffusion, whereas 61% (T-T) and 67% (H-T) showed free diffusion (**Figure 3B**). We next determined the jump angle for the different segments in the single-molecule trajectories (see Methods), which represents an additional measure of local confinement of protein mobility.[45,46] Consistently, we found a more confined pattern for the FRET segment (dimeric receptor complex) as compared to a random walk pattern for the donor-only segment (monomeric receptor) (**Figure 3C**). We also examined the diffusion coefficient and mode of MET dimers for trajectories that began with a FRET signal and transitioned to a donor-only signal after the acceptor signal disappeared (pathway 1) as compared to those that showed a loss in FRET signal without remaining donor-only signal (pathway 2) (**Figure S3**). While the diffusion coefficient of the FRET segment was found to be similar in both pathways (0.066 ± 0.044 µm^2^/s and 0.060 0.043 µm^2^/s in T-T labeled cells; 0.056 ± 0.038 µm^2^/s and 0.055 ± 0.040 µm^2^/s in H-T labeled cells) (**Figure S3B**), an increase in the fraction with restricted diffusion was observed (from 44% (pathway 1) to 60% (pathway 2) for T-T labelled cells, and from 50% (pathway 1) to 59% (pathway 2) in H–T labelled cells) (**Figure S3C**). The jump angle distribution for these segments of trajectories showed a similarly restricted motion (**Figure S3D**).

**Figure 3:**
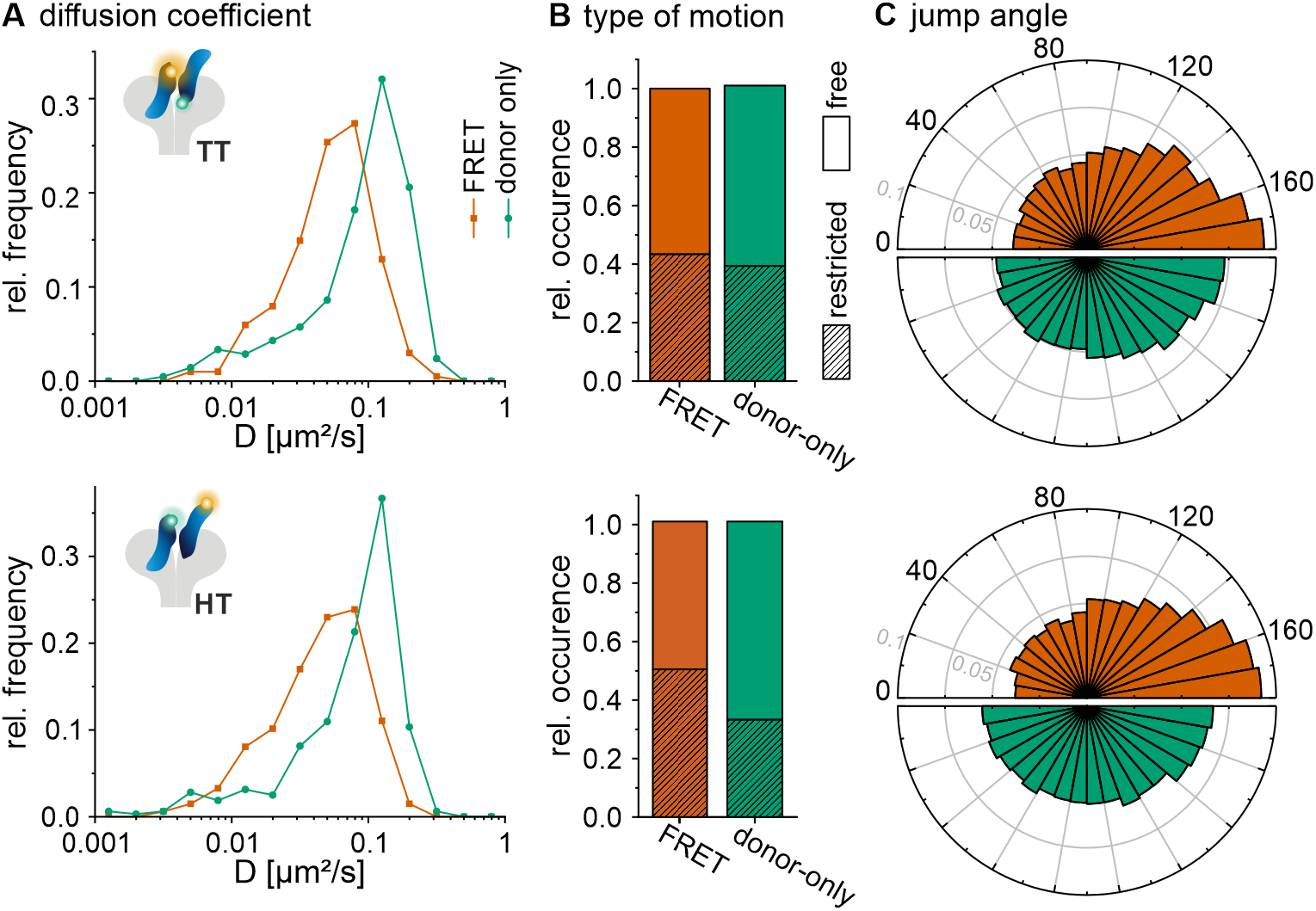
Diffusion dynamics of MET:InlB extracted from FRET trajectories. **A**) Distribution of diffusion coefficients for the FRET segment (orange) and the donor-only segment (green) of single-molecule trajectories (N = 27 cells, 24 cells measured with InlB T-T variants and H-T variants, respectively, from at least 3 independent experiments). **B**) Relative occurrence of restricted and free diffusion MET:InlB within FRET and donor-only segments of single-molecule trajectories. **C**) Distribution of jump angles within FRET and donor-only segments of single-molecule trajectories (grey circles represent relative frequencies).

## Discussion

The probability of detecting a single-molecule FRET event depends on the degree of labeling. For high protein expression levels, as observed for MET in many relevant cell lines,^[29]^ stoichiometric labeling may, however, preclude single-molecule detection. SPT in particular demands molecular densities smaller than the diffraction limit, ideally below 0.5 receptors/µm^2^.^[47]^ To overcome these challenges, smFRET combined with fluorescence recovery after photobleaching (RAP) has proven particularly effective for studying native membrane receptors.^[40]^ RAP minimizes signal overlap and background fluorescence, improving the specificity of single-molecule detection while maintaining a high labeling efficiency. We combined smFRET-RAP with single-particle tracking to detect dimer–monomer transitions of MET receptors in living cells and with single-molecule resolution. In addition, smFRET-RAP reduces fluorescence resulting from fluorophore-labeled ligands nonspecifically adsorbed to the surface. For instance, the hydrophobic ATTO 647N fluorophore used in this study is known to show increased surface binding, which in the SPT analysis appears as an immobile fraction and leads to an underestimation of diffusion coefficients.^[48,49]^ These artifacts were minimized by initial photobleaching in the smFRET-RAP workflow before the measurement. In addition, selecting only FRET trajectories with anti-correlated donor and acceptor signals ensures high specificity in the analyzed single-molecule trajectories. Consistently, global mean squared displacement (MSD) analysis of FRET-positive trajectories revealed negligible immobile diffusion, validating the robustness of the approach.

The distributions of FRET efficiencies of the (MET:InlB_321_)_2_ dimer recorded in living cells were broader than those in fixed cells (**Figure 1C, S2A**). Several parameters contribute to this observation: first, single-particle tracking in living cells requires shorter integration times to reliably capture the mobility of single molecules in the cell membrane.^[47,50,51]^ Related to this, lower laser intensities were used in live-cell experiments to minimize phototoxicity, which in turn impacts signal-to-noise and signal-to-background values. Second, background intensities in live-cell experiments may be different and vary during an experiment. Lastly, it is possible that the (MET:InlB_321_)_2_ complex exhibits some degree of structural flexibility in living cells, which, however, is beyond the current accuracy of the method.

The combination of single-molecule FRET and single-particle tracking allows for a segmentation of single-molecule trajectories, and thus a separate analysis of differently built molecular complexes. Single-molecule FRET trajectories of the (MET:InlB_321_)_2_ dimer showed a loss in acceptor fluorescence after an average of 1.3 (InlB T-T) or 0.8 (InlB H-T) seconds (**Figure 2C**), whereas smFRET experiments in fixed cells showed a much longer duration of the FRET signal (of 19 and 13 seconds, respectively). This indicates that the duration of the FRET signal in live-cell experiments is predominantly determined by the lifetime of the (MET:InlB_321_)_2_ dimer, and not influenced by photobleaching of the acceptor fluorophore. The loss in acceptor fluorescence in live-cell experiments can be further classified into two reaction pathways: continuous donor fluorescence in the single-molecule trajectory indicates at the dissociation of the (MET:InlB_321_)_2_ dimer, whereas the simultaneous disappearance of both the donor and acceptor signals indicates that the dimeric complex moved beyond the TIRF detection limit,^[40,52]^ possibly through cellular uptake such as e.g. endocytosis ^[15,53]^. Other explanations are possible, such as the off-dissociation of InlB ligands from the dimeric complex, yet are not accessible from the experimental data. We note that this study entirely focuses on the lifetime of (MET:InlB)_2_ dimers. Previous work has shown that MET also forms receptor dimers in the absence of ligands.^[25,26]^

The determined lifetime of the (MET:InlB_321_)_2_ dimer is similar to previously reported values for EGFR:EGF dimers ^[54–56]^. Using a similar fluorophore-based labeling approach, Coban et al. reported an (EGFR:EGF)_2_ complex lifetime of approximately 0.84 s in the breast cancer cell line HCC1054.^[54]^ Another study that employed quantum dot (QD)-labeled EGF in combination with hidden Markov modeling revealed longer lifetimes of EGFR:EGF dimers and a variation across different cell lines, with 3.59 s in the epidermoid carcinoma cell line A431, 13.02 s in HeLa cells, and 8.33 s in EGFR-transfected CHO cells.^[55,56]^ The longer lifetimes might be related to the larger QD labels, which exhibit outstanding brightness yet might also impede receptor mobility and subsequent processing steps of ligand-activated receptors. Notably, the diffusion coefficient of EGFR:EGF varied from 0.07 µm^2^/s with eGFP labeling to 0.014 µm^2^/s with QD labeling in CHO cells.^[56,57]^

From segmented trajectories, we observed a reduced diffusion coefficient of the dimeric (MET:InlB)_2_ complex in the FRET segment as compared to the monomeric (dissociated) MET:InlB complex from the donor-only segment (**Figure 3**). Previous SPT studies have reported a reduction of the global diffusion coefficients of RTKs, including MET, upon ligand stimulation.^[30,50,51,58–60]^ In this work, we report the measurement of diffusion coefficients of the same receptor-ligand complex for two distinct molecular stoichiometries.

An alternative technology to investigate protein-protein interactions in the plasma membrane of living cells is two-color single-particle tracking (2cSPT).^[47,55,61–66]^ In 2cSPT experiments, dimers are identified by assigning two particles in two spectrally separate channels as co-localized if the distance is below a defined value.^[47,61,65–67]^ The upper threshold for this distance threshold is determined by the localization precision of each channel and the accuracy of channel alignment.^[61]^ However, in studies involving RTKs, which are mainly internalized through clathrin-mediated endocytosis (CME),^[16,68,69]^ it is challenging to discriminate between RTK dimers and RTKs colocalized within clathrin-coated pits that exhibit a size from 70 to 150 nm and persist for more than tens of seconds.^[70–73]^ In contrast, smFRET in living cells offers an elegant alternative to detect *bona fide* receptor dimers, given the short range of the FRET interaction of less than 10 nm and the feature of anti-correlated donor and acceptor intensity traces._[40,47,67]_

Following ligand binding and receptor dimerization, intracellular signaling is initiated by adaptor protein recruitment, such as Grb2 binding to ligand-activated MET. Previous studies reported that Grb2 binds to activated EGFR with a lifetime of approximately 0.13 s in both *in vivo* and *in vitro* systems.^[74,75]^ Within this context, the ∼1-second lifetime of (MET:InlB)_2_ dimers seems plausible. Following signaling initiation, RTKs may also be recycled through CME, a key regulatory mechanism for RTK downregulation. CME involves an initial approximately 10 second phase for clathrin-coated pit formation, followed by membrane invagination and vesicle internalization over 20 seconds to several minutes.^[71,72]^

In summary, the presented experimental approach presents a robust method to measure the stability of ligand-activated membrane receptors in living cells, and to assess the diffusion properties of differently constituted receptor complexes. The approach is transferable to other membrane proteins, and it can serve as a platform to assess how modifications in proteins or the addition of modulators alter the stability of receptor dimers.

## Author contributions

MH and YL designed the research. HHN designed the FRET study and provided labeled proteins. YL performed live-cell single-molecule imaging experiments and image analysis with support from MSD. HDB built the optical setup for smFRET. YL, HHN, MSD, and MH discussed the data and wrote the manuscript.

## Acknowledgments

We are grateful to Inga Hänelt and Roberto Covino for helpful discussions. We thank Petra Freund for support with cell culture and Wenjun Wang for assistance with coding.

## Data availability

The data is available in the EMBL BioImaging Archive under accession code S-BIAD1347 (https://www.ebi.ac.uk/biostudies/bioimages/studies/S-BIAD1347). Custom-written software used in this work was written in Python and MatLab and is available from the authors upon request.

