## supplemental information for "Single-Molecule FRET-Tracking of InlB-Activated MET Receptors in Living Cells"

##### Supplementary Movies

**Supplementary Movie 1:** Single-molecule FRET movie of MET receptors labeled with InlB-T-Cy3B and InlB-T-ATTO 647N. Scale bar 1 µm. Corresponding to Figure S1A, upper panel.

**Supplementary Movie 2:** Single-molecule FRET movie of MET receptors labeled with InlB-T-Cy3B and InlB-T-ATTO 647N. Scale bar 1 µm. Corresponding to Figure S1A, bottom panel.

**Supplementary Movie 3:** Single-molecule FRET movie of MET receptors labeled with InlB-H-Cy3B and InlB-T-ATTO 647N. Scale bar 1 µm. Corresponding to Figure S1B, upper panel.

**Supplementary Movie 4:** Single-molecule FRET movie of MET receptors labeled with InlB-H-Cy3B and InlB-T-ATTO 647N. Scale bar 1 µm. Corresponding to Figure S1B, bottom panel.

### Supplementary Figures

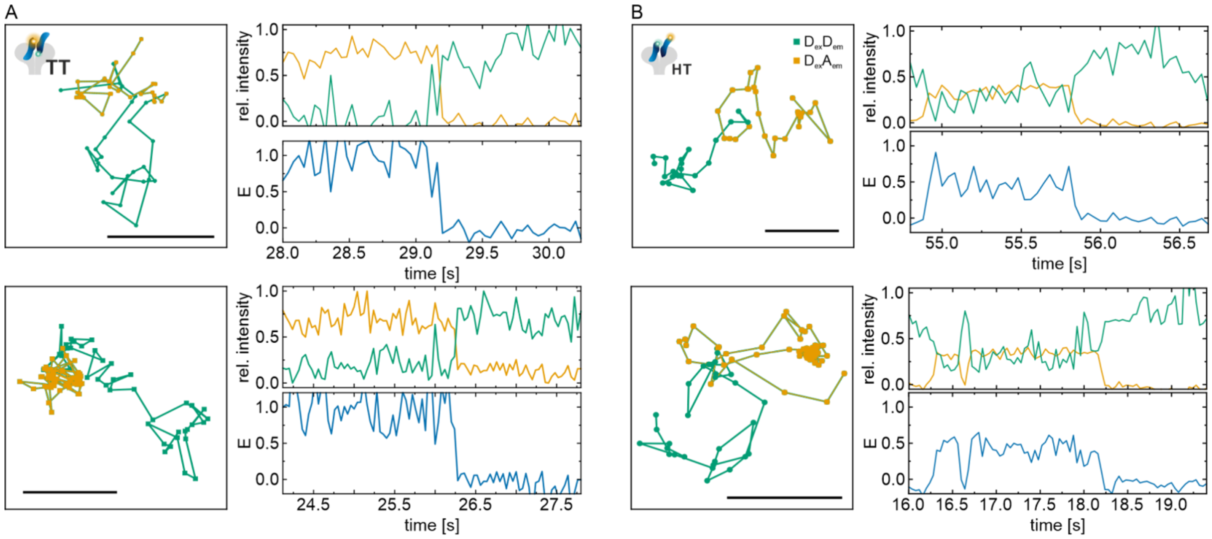


**Figure S1: smFRET-RAP of (MET:InlB$_{321}$)$_2$ dimers in living U-2 OS cells.** Exemplary smFRET trajectories for **A**) Cy3B-T-InlB$_{321}$ and ATTO 647N-T-InlB$_{321}$ and **B**) Cy3B-H-InlB$_{321}$ and ATTO 647N-T-InlB$_{321}$ (right). Trajectories are shown with the donor (green) and acceptor (orange) intensity traces upon donor excitation and the respective FRET efficiencies (blue). Scale bars are 500 nm.

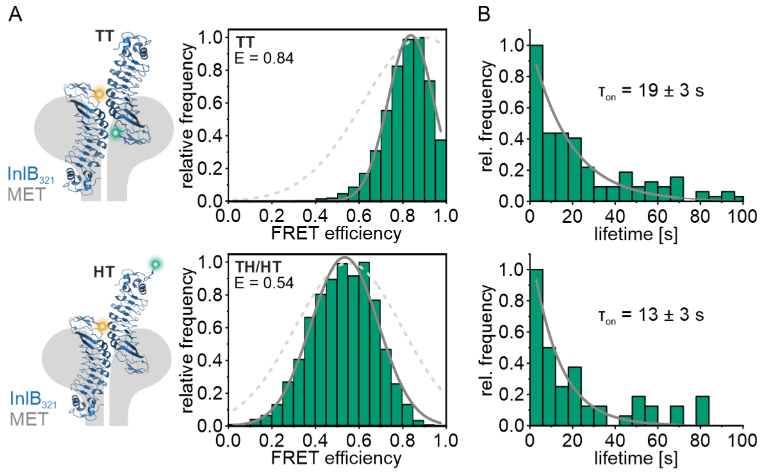


**Figure S2: smFRET of $(MET{:}InlB_{321})_2$ dimers in fixed cells. A**) FRET efficiency distributions for Cy3B-T-$InlB_{321}$/ATTO 647N-T-$InlB_{321}$ (N = 113 smFRET traces from 64 cells) and Cy3B-H-$InlB_{321}$/ATTO 647N-T-$InlB_{321}$ (N = 49 smFRET traces from 39 cells) are displayed. Overlay is the FRET efficiency distribution of living cells (light gray dotted line). **B**) The corresponding lifetimes of smFRET traces were histogrammed and fitted with a single-exponential decay.

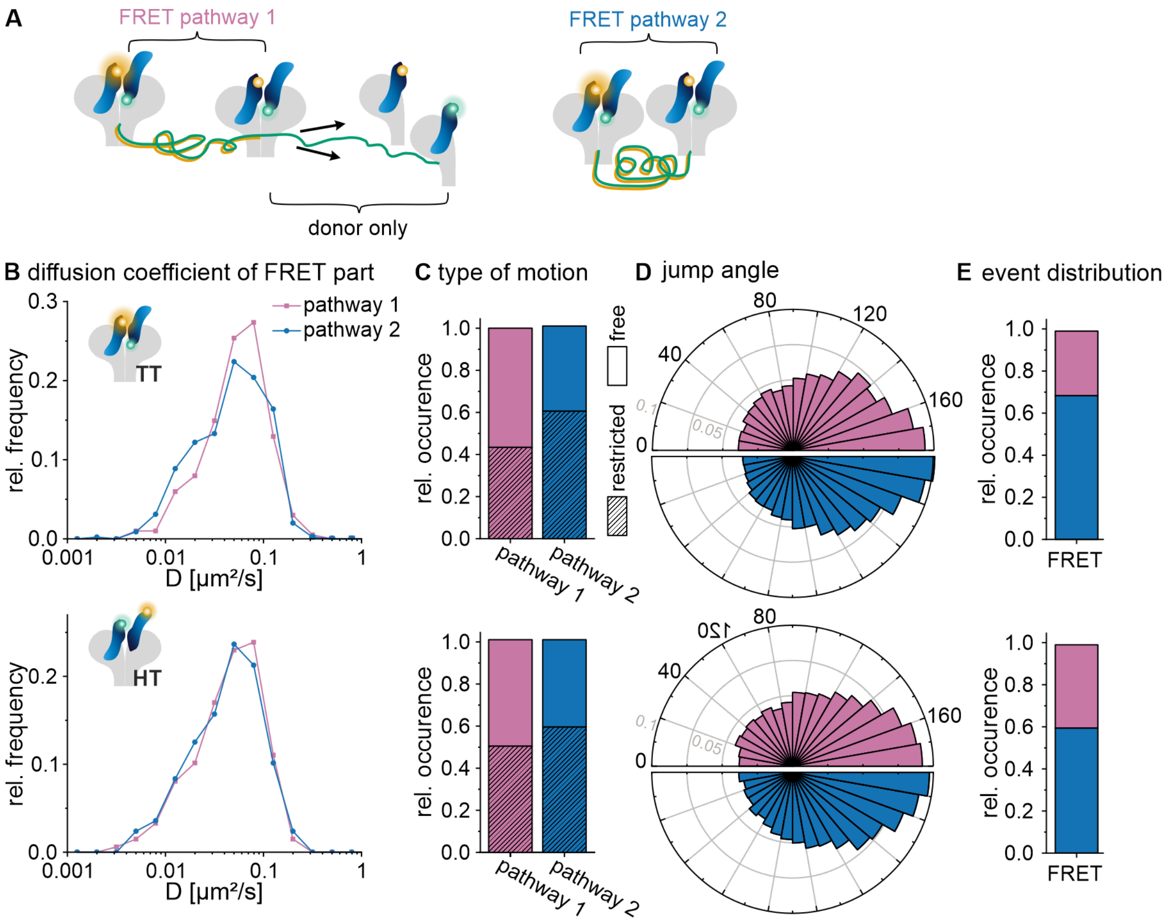


**Figure S3: Dynamics of (MET:InlB$_{321}$)$_2$ dimer detected with smFRET in two pathways. A**) Schematic of two possible pathways observed in single-molecule FRET trajectories. **B**) Distribution of diffusion coefficients determined for the FRET segments in single-molecule trajectories for pathway 1 (pink) and pathway 2 (blue) (N = 27 cells (InlB T-T) and 24 cells (InlB H-T), respectively, and from at least 3 independent experiments). **C**) Relative occurrence of restricted and free diffusion in FRET segments of pathway 1 and 2. **D**) Distribution of jump angles of FRET segments of single-molecule trajectories in pathways 1 and 2 (grey scales represent relative frequencies). **E**) Relative occurrence of pathway 1 and pathway 2 in single-molecule trajectories.

#### Supplementary Table

**Table S1: Degree of labeling of fluorophore-labeled InlB$_{321}$ variants.** InlB variants were labeled either with Cy3B or ATTO 647N maleimide. The cysteine mutations used for fluorophore labeling are indicated. The degree of labeling (DOL) was determined by absorption spectroscopy.

| Variant | Mutation | DOL (%) |
|---|---|---|
| InlB-T-Cy3B | K280C | 87 |
| InlB-T-ATTO 647N | K280C | 69 |
| InlB-H-Cy3B | K64C | 70 |
| InlB-H-ATTO 647N | K64C | 103 |
